# MaveDB v2: a curated community database with over three million variant effects from multiplexed functional assays

**DOI:** 10.1101/2021.11.29.470445

**Authors:** Alan F Rubin, Joseph K Min, Nathan J Rollins, Estelle Y Da, Daniel Esposito, Matthew Harrington, Jeremy Stone, Aisha Haley Bianchi, Mafalda Dias, Jonathan Frazer, Yunfan Fu, Molly Gallaher, Iris Li, Olivia Moscatelli, Jesslyn YL Ong, Joshua E Rollins, Matthew J Wakefield, Shenyi “Sunny” Ye, Amy Tam, Abbye E McEwen, Lea M Starita, Vanessa L Bryant, Debora S Marks, Douglas M Fowler

## Abstract

A central problem in genomics is understanding the effect of individual DNA variants. Multiplexed Assays of Variant Effect (MAVEs) can help address this challenge by measuring all possible single nucleotide variant effects in a gene or regulatory sequence simultaneously. Here we describe MaveDB v2, which has become the database of record for MAVEs. MaveDB now contains a large fraction of published studies, comprising over two hundred datasets and three million variant effect measurements. We created tools and APIs to streamline data submission and access, transforming MaveDB into a hub for the analysis and dissemination of these impactful datasets.

## Background

Genomes contain both genes that encode RNAs and proteins and non-coding regulatory elements that modulate gene expression. Variation within genomes produces interindividual differences that govern a multitude of traits, including many that relate to disease. Because DNA sequencing is now inexpensive and widely deployed, human genetic variants are being discovered at a staggering pace. For example, approximately 241 million small variants comprising single nucleotide changes and small deletions/insertions have been identified in 140,000 individuals in gnomAD [1]. Of these variants, 4.6 million are single amino acid changes (i.e. missense variants). 4.5 million missense variants have been identified among 200,000 individuals in the UK Biobank [2]. In contrast, only 437,000 missense variants have been annotated in ClinVar, of which greater than 70% are currently variants of uncertain significance and thus cannot be used for clinical decision making [3]. Understanding how these existing variants, as well as the ones we will discover as more individuals are sequenced, impact molecular, cellular and organismal phenotypes represents a central challenge for genomics.

In the past, genetic variants would be tested for functional effects in bespoke assays singly or in relatively low numbers, but more recent technologies have enabled the increasingly popular multiplexed assays of variant effect (MAVEs) approaches [4]. In a MAVE, the functional effects of thousands or tens of thousands of variants of a DNA regulatory region, coding genes, UTRs or other functional elements are simultaneously experimentally determined. To achieve this scale, a large library of variants is made and tested in a pooled fashion, with high-throughput DNA sequencing used to read out variant effects (for a detailed description see [5–7]). The result is a comprehensive variant effect map, which contains the experimentally measured effects of most or all of the possible single nucleotide or missense variants. Variant effect maps are of high utility. For example, in genes where germline variants can increase disease risk, variant effect maps can help to resolve up to ∼70% of clinical variants of uncertain significance [8]. Variant effect maps can also be used to probe the protein sequence/function relationship [9–42], assist in protein design [43], reveal protein structure [44,45], elucidate regulatory DNA and gene function [46–49] and train or evaluate variant effect predictors [50–52]. Thus, multiplexed functional data are poised to have a major impact, and efforts are now underway to scale up and apply MAVEs to a significant fraction of the human genome [53–55].

However, multiplexed functional data availability has been a major hurdle. In 2019 we created MaveDB, a public, open source repository for multiplexed functional data [56]. MaveDB allows researchers to store, share, and access processed multiplexed variant functional data and associated metadata in a standardized, searchable format. MaveDB implements an easy-to-use web interface that facilitates data deposition by researchers. However, MaveDB suffered from three key limitations. First, it contained only a small fraction of the data available at the time. Second, data from new multiplexed assays such as saturation genome editing [14,57] were not compatible with the original MaveDB data model. Finally, MaveDB was not designed with federation across genomic data resources in mind.

Here we describe several technical advances and data model improvements implemented in MaveDB v2, as well as provide a progress update on our extensive curation of previously-published multiplexed assay results and expansion of database content. We have improved the user experience for contributors by adding API-based user uploads and a companion Python module for researchers with bioinformatics or data science experience. We have updated our data model by adding a new type of record for imputation or the combination of results across multiple experiments. We have also refined and formalized our variant representation, allowing us to support more diverse types of MAVE variants and associated experimental designs, while also improving compatibility with existing international standards. In addition to these technical improvements, we have curated 161 new datasets from the literature, encompassing two million new variants and more than tripling the total number of variant effect measurements in the database.

## Database functionality and content

MaveDB is designed to store and distribute multiplexed variant functional data, including scores, associated data and metadata. A MAVE dataset minimally includes a collection of variant effect scores that describe the functional consequences of nucleotide or amino acid variants. Each variant effect score can also be associated with a variance estimate, variant frequency or other data. The metadata includes descriptions of the experimental and data analysis methods and target information, such as the sequence and accession numbers in other databases.

MaveDB has a hierarchical structure populated by score set, experiment, and experiment set records. Score sets contain the variant effect scores and associated data columns, such as variance estimates and variant counts, details about the experimental target sequence, and a description of the score calculations. Experiments group together multiple score sets that depend on the same raw data, thereby improving discoverability for users, and describe the assay that was performed. Experiment sets do not have any data or metadata themselves, but group related experiments, such as multiple assays performed on a single target and described in the same publication.

### Meta-analysis score sets

In addition to standard score sets linked to an experiment and its raw data, MaveDB v2 implements meta-analysis score sets. These new records are linked to one or more existing score set records, and are used to describe a distinct version of the scores that have been transformed in some way. For example, a dataset that has imputed the values of missing scores would be best represented as a meta-analysis score set linked to the pre-imputation scores, ensuring the pre-imputation scores are preserved and discoverable. Another example is a set of scores that result from the combination of multiple existing score sets (Figure 1).

**Figure 1:**
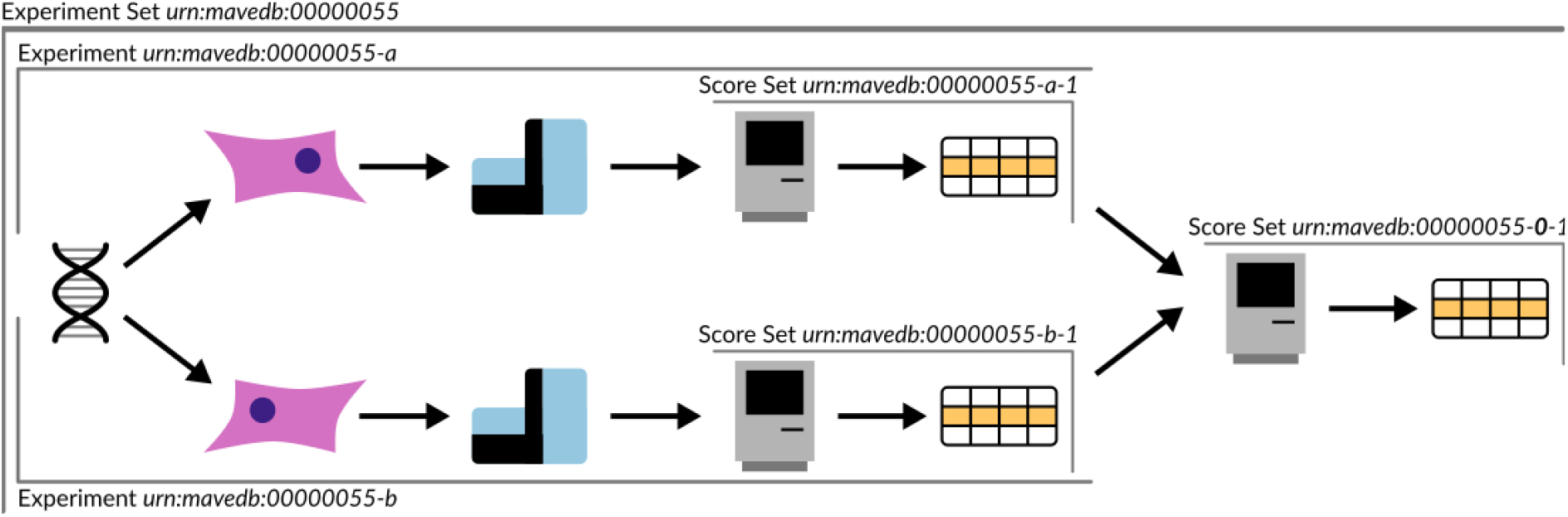
Representative structure for a meta-analysis score set. The cartoon depicts the relationship between experiment sets, experiments, score sets, and meta-analysis score sets for a real-world dataset. Two assays were performed on the gene *NUDT15* and combined into a resulting “function score” that summarized performance across both assays [41]. The meta-analysis score set that combines these results is shown on the far right, and is described in MaveDB under urn:mavedb:00000055-0-1.

We recommend that users upload minimally-transformed scores as standard score sets to MaveDB, and create new meta-analysis score sets that describe normalization or imputation steps as applicable. This enables other researchers who want to build models that would be sensitive to data normalization or evaluate their own normalization methods to make use of the data without having to perform a full re-analysis from counts or sequence reads.

With its hierarchical data model and the introduction of meta-analysis score sets, MaveDB enables provenance tracking for individual variant measurements from a multiplexed assay. This tracking is critical for downstream usage, particularly in clinical pipelines, where it is essential to understand the independence of different data sources for evaluating variant pathogenicity using the ACMG guidelines [58].

### Standardized variant representation

Previously MaveDB used a variant representation based on the Enrich2 [59] output format, which was in turn inspired by the HGVS Sequence Variant Nomenclature [60]. Starting with MaveDB v2, we have adopted a revised variant representation we call MAVE-HGVS. MAVE-HGVS is a subset of HGVS version 20.05. HGVS nomenclature is comprehensive and very expressive, and consequently includes syntax that is not needed to represent variants from multiplexed variant functional data. While packages exist for parsing HGVS [61,62], they are intended for use in human genetics and rely on sequence database entries that are not always available for multiplexed assays. MAVE-HGVS is an easy-to-parse subset of the HGVS nomenclature that captures the types of variants that routinely occur in MAVE datasets while not relying on external sequence databases or identifiers.

In addition to the specification, MAVE-HGVS has a reference Python implementation, mavehgvs. We use this implementation in MaveDB v2 to validate variants when multiplexed variant functional data are uploaded to the database, both to make sure that the variant strings are in a valid format and to ensure that variants are consistent with the target sequence.

MAVE-HGVS offers three major improvements over the old MaveDB format. First, MAVE-HGVS is much more easily convertible to standard HGVS. Second, the previous format did not define target-identical (“wild type”) variants in a standard way and was not able to define target-identical variants by position, a feature that is needed for MITE-Seq datasets [31]. Finally, we can continue to draw from HGVS as support for new variant types is needed. We have already taken advantage of this by adding splice variants to more faithfully represent datasets curated from the literature.

### Community tools

To empower the MAVE community, we developed MaveTools, a Python library designed for interactive development environments like Jupyter notebooks [63], replacing our previous command-line tool mavedbconvert. MaveTools has functions to convert commonly-used variant representations such as Enrich seqid [64] to MAVE-HGVS and create MAVE-HGVS strings that describe the differences between pairs of codons. MaveTools also implements a local version of the dataset validation logic used by the MaveDB server, making it easier for users to identify and resolve formatting errors. In addition to MaveDB-specific features, we are augmenting the library with general utility functions for researchers working with MAVE datasets.

MaveDB, mavehgvs, and MaveTools are all distributed under OSI-approved open source licenses and development activity is ongoing in their public GitHub repositories (see Code availability section). We envision that MaveTools will become a valuable community resource, particularly for students and others who are new to working with MAVE data. We welcome contributions and engagement from other members of the community in those spaces or by contacting us directly.

### Improved API support

The previous version of MaveDB only accepted data via web form. We now also support data deposition through REST API. The API uses the same logic and validation as the web interface, ensuring continuity and data integrity. This makes depositing some datasets much more efficient, such as a series of similar assays that measure variant effects under different concentrations of a small molecule. We hope that authors of MAVE analysis pipelines will consider adopting this API endpoint as an output option.

MaveTools provides REST API wrappers so users with a bioinformatics or data science background can query the database and upload or download datasets. By formatting a Python object that contains appropriate metadata and file paths for score and count files, users can validate the data and upload it to the server within a script or interactive session. Researchers who are building machine learning models or otherwise using large amounts of MAVE data can also use the wrappers to automate downloads. The MaveTools documentation and source code repository includes example Jupyter notebooks for both upload and download that users can follow and modify.

We have also added a new API endpoint that allows users to download data on individual variants. This feature currently only supports the MaveDB variant IDs that are assigned when the variants are created in our database, but we are in the process of mapping these IDs to more widely-used accession numbers, particularly for genes of clinical relevance.

### Database curation and contents

When the original MaveDB manuscript was published in 2019, data from only 7% of publications at the time were included. Since then, as with many databases, the gap in representation has grown. Thus, we launched a concerted effort to deposit published datasets that were not yet included in MaveDB. Our curation team spanned three sites: WEHI/University of Melbourne in Melbourne, Australia; University of Washington in Seattle, USA; and Harvard University in Boston, USA.

To enable the discovery, reuse, and interpretation of data in MaveDB, we developed a robust process for curating and summarizing these heterogeneous experimental results, including training materials, much of which has been incorporated in the updated documentation pages in MaveDB v2. Key information was abstracted from the associated publications and synthesised into a title, short description, abstract, and methods for each experiment and score set record. Accession numbers for raw data and target sequences for each dataset were also included. Each draft MaveDB entry was peer reviewed by at least one other MaveDB curator to ensure all relevant information was present and accurate before inclusion in the database. In addition to writing the free text sections and organizing associated metadata, our curation team also formatted scores and related values from supplemental tables.

As a result of these efforts, MaveDB now contains 231 datasets from 68 publications. This comprises 46% of all published papers for which data was available, 28% of published papers overall (Figure 2). Furthermore, MaveDB now makes available 20 datasets from 7 papers that were not provided as part of their original publications.

**Figure 2:**
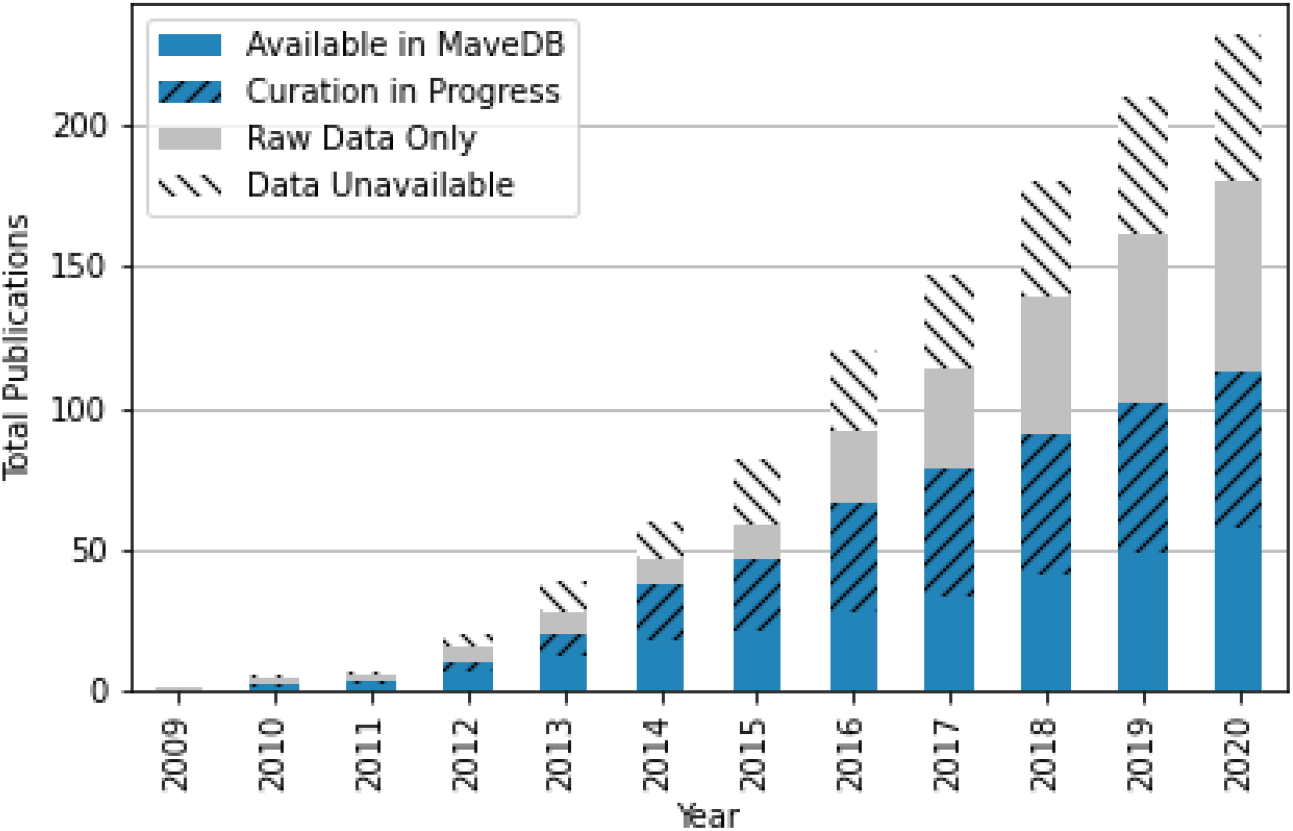
Growth of datasets and availability over time. We compiled a list of 238 publications that contained at least one new MAVE dataset and determined whether counts, scores, or raw sequence data were made available in supplementary information. At present, we have made 46% of all datasets for which scores were available accessible via MaveDB, and work is underway to add the remaining datasets.

## Conclusions

Measuring, predicting and understanding variant effects on a genome-wide scale are key challenges. MAVEs are an important approach for overcoming these challenges, but the data must be stored in a stable, standardized fashion along with the metadata required to enable reanalysis, modeling and prediction. Moreover, MAVE datasets must be easily discoverable, and MAVE data must appear in other commonly used data portals.

MaveDB v2 marks a major improvement to our data model and ability to handle these heterogeneous datasets. It improves upon the previously-used variant representation and furthers our aim to standardize and share data. MaveDB v2 also incorporates new API features, including the ability to request data on individual variants and upload data via script. These features are supported by open source software designed to serve the research community.

Additionally, we launched a massive curation effort involving 161 datasets, ultimately populating MaveDB with nearly half of all published data. This equates to three million variant measurements. Going forward we strongly encourage data generators to deposit their data directly into MaveDB prior to or upon publication.

## Code availability

MaveDB source code is available at https://github.com/VariantEffect/MaveDB. It is distributed under the AGPLv3 software license.

mavehgvs source code is available at https://github.com/VariantEffect/mavehgvs. It is distributed under the 3-Clause BSD software license.

MaveTools source code is available at https://github.com/VariantEffect/MaveTools. It is distributed under the 3-Clause BSD software license.

## Acknowledgements

The Atlas of Variant Effects (AVE) Alliance Data Coordination and Dissemination workstream contributed valuable feedback on the design and goals of MaveDB. This work was supported by the National Institutes of Health (NIH; RM1HG010461 to DMF, UM1HG011969 to LMS and DMF) and by Chan Zuckerberg Initiative (CZI2018-191853 to DSM) The research benefited from support from the Victorian State Government Operational Infrastructure Support and Australian Government NHMRC Independent Research Institute Infrastructure Support.

## Author contributions

AFR and DMF designed the database. AFR JKM NJR EYD DE MH and JS wrote the database and associated software. AFR NJR MH AHB MD JF YF MG IL OM JYLO JER MJW SY AT AEM and DSM curated datasets. AFR VLB DSM and DMF supervised dataset curation. AFR LMS DSM and DMF supervised the software projects. AFR AHB OM SY and DMF wrote the paper.

## References

1. Karczewski KJ, Francioli LC, Tiao G, Cummings BB, Alföldi J, Wang Q, et al. The mutational constraint spectrum quantified from variation in 141,456 humans. Nature. 2020;581:434–43.

2. Bycroft C, Freeman C, Petkova D, Band G, Elliott LT, Sharp K, et al. The UK Biobank resource with deep phenotyping and genomic data. Nature. 2018;562:203–9.

3. Landrum MJ, Lee JM, Benson M, Brown GR, Chao C, Chitipiralla S, et al. ClinVar: improving access to variant interpretations and supporting evidence. Nucleic Acids Res. 2018;46:D1062–7.

4. Starita LM, Ahituv N, Dunham MJ, Kitzman JO, Roth FP, Seelig G, et al. Variant Interpretation: Functional Assays to the Rescue. The American Journal of Human Genetics. 2017;101:315–25.

5. Fowler DM, Fields S. Deep mutational scanning: a new style of protein science. Nat Meth. 2014;11:801–7.

6. Kinney JB, McCandlish DM. Massively Parallel Assays and Quantitative Sequence–Function Relationships. Annual Review of Genomics and Human Genetics. 2019;20:null.

7. Weile J, Roth FP. Multiplexed assays of variant effects contribute to a growing genotype-phenotype atlas. Hum Genet. 2018;137:665–78.

8. Fayer S, Horton C, Dines JN, Rubin AF, Richardson ME, McGoldrick K, et al. Closing the gap: Systematic integration of multiplexed functional data resolves variants of uncertain significance in BRCA1, TP53, and PTEN. Am J Hum Genet. 2021;S0002-9297(21)00411-0.

9. Ashenberg O, Padmakumar J, Doud MB, Bloom JD. Deep mutational scanning identifies sites in influenza nucleoprotein that affect viral inhibition by MxA. PLoS Pathog. 2017;13:e1006288.

10. Bandaru P, Shah NH, Bhattacharyya M, Barton JP, Kondo Y, Cofsky JC, et al. Deconstruction of the Ras switching cycle through saturation mutagenesis. Valencia A, editor. eLife. 2017;6:e27810.

11. Chen JZ, Fowler DM, Tokuriki N. Comprehensive exploration of the translocation, stability and substrate recognition requirements in VIM-2 lactamase. EdFleishman SJ, Marletta MA, Ostermeier M, Whitehead TA, editors. eLife. eLife Sciences Publications, Ltd; 2020;9:e56707.

12. Chiasson MA, Rollins NJ, Stephany JJ, Sitko KA, Matreyek KA, Verby M, et al. Multiplexed measurement of variant abundance and activity reveals VKOR topology, active site and human variant impact. Elife. 2020;9:e58026.

13. Doud MB, Bloom JD. Accurate Measurement of the Effects of All Amino-Acid Mutations on Influenza Hemagglutinin. Viruses. 2016;8:155.

14. Findlay GM, Daza RM, Martin B, Zhang MD, Leith AP, Gasperini M, et al. Accurate classification of BRCA1 variants with saturation genome editing. Nature. Nature Publishing Group; 2018;562:217–22.

15. Firnberg E, Labonte JW, Gray JJ, Ostermeier M. A Comprehensive, High-Resolution Map of a Gene’s Fitness Landscape. Mol Biol Evol. 2014;31:1581–92.

16. Fowler DM, Araya CL, Fleishman SJ, Kellogg EH, Stephany JJ, Baker D, et al. High-resolution mapping of protein sequence-function relationships. Nat Meth. 2010;7:741–6.

17. Giacomelli AO, Yang X, Lintner RE, McFarland JM, Duby M, Kim J, et al. Mutational processes shape the landscape of TP53 mutations in human cancer. Nature Genetics. 2018;1.

18. Glazer AM, Kroncke BM, Matreyek KA, Yang T, Wada Y, Shields T, et al. Deep Mutational Scan of an SCN5A Voltage Sensor. Circ Genom Precis Med. 2020;13:e002786.

19. Gray VE, Sitko K, Kameni FZN, Williamson M, Stephany JJ, Hasle N, et al. Elucidating the Molecular Determinants of Aβ Aggregation with Deep Mutational Scanning. G3 (Bethesda). 2019;9:3683–9.

20. Jia X, Burugula BB, Chen V, Lemons RM, Jayakody S, Maksutova M, et al. Massively parallel functional testing of MSH2 missense variants conferring Lynch syndrome risk. Am J Hum Genet. 2021;108:163–75.

21. Jones EM, Lubock NB, Venkatakrishnan AJ, Wang J, Tseng AM, Paggi JM, et al. Structural and functional characterization of G protein-coupled receptors with deep mutational scanning. Elife. 2020;9:e54895.

22. Kitzman JO, Starita LM, Lo RS, Fields S, Shendure J. Massively parallel single-amino-acid mutagenesis. Nat Meth. 2015;12:203–6.

23. Kotler E, Shani O, Goldfeld G, Lotan-Pompan M, Tarcic O, Gershoni A, et al. A Systematic p53 Mutation Library Links Differential Functional Impact to Cancer Mutation Pattern and Evolutionary Conservation. Molecular Cell. 2018;71:178–190.e8.

24. Lee JM, Huddleston J, Doud MB, Hooper KA, Wu NC, Bedford T, et al. Deep mutational scanning of hemagglutinin helps predict evolutionary fates of human H3N2 influenza variants. PNAS. 2018;201806133.

25. Majithia AR, Tsuda B, Agostini M, Gnanapradeepan K, Rice R, Peloso G, et al. Prospective functional classification of all possible missense variants in PPARG. Nat Genet. 2016;48:1570–5.

26. Matreyek KA, Starita LM, Stephany JJ, Martin B, Chiasson MA, Gray VE, et al. Multiplex assessment of protein variant abundance by massively parallel sequencing. Nature Genetics. 2018;50:874–82.

27. Mavor D, Barlow K, Thompson S, Barad BA, Bonny AR, Cario CL, et al. Determination of ubiquitin fitness landscapes under different chemical stresses in a classroom setting. eLife Sciences. 2016;5:e15802.

28. McLaughlin Jr RN, Poelwijk FJ, Raman A, Gosal WS, Ranganathan R. The spatial architecture of protein function and adaptation. Nature. 2012;491:138–42.

29. Mehlhoff JD, Stearns FW, Rohm D, Wang B, Tsou E-Y, Dutta N, et al. Collateral fitness effects of mutations. PNAS. National Academy of Sciences; 2020;117:11597–607.

30. Melamed D, Young DL, Gamble CE, Miller CR, Fields S. Deep mutational scanning of an RRM domain of the Saccharomyces cerevisiae poly(A)-binding protein. RNA. 2013;19:1537–51.

31. Melnikov A, Rogov P, Wang L, Gnirke A, Mikkelsen TS. Comprehensive mutational scanning of a kinase in vivo reveals substrate-dependent fitness landscapes. Nucleic Acids Res. 2014;42:e112–e112.

32. Mighell TL, Evans-Dutson S, O’Roak BJ. A Saturation Mutagenesis Approach to Understanding PTEN Lipid Phosphatase Activity and Genotype-Phenotype Relationships. The American Journal of Human Genetics. 2018;102:943–55.

33. Mishra P, Flynn JM, Starr TN, Bolon DNA. Systematic Mutant Analyses Elucidate General and Client-Specific Aspects of Hsp90 Function. Cell Reports. 2016;15:588–98.

34. Ogden PJ, Kelsic ED, Sinai S, Church GM. Comprehensive AAV capsid fitness landscape reveals a viral gene and enables machine-guided design. Science. 2019;366:1139–43.

35. Romero PA, Tran TM, Abate AR. Dissecting enzyme function with microfiuidic-based deep mutational scanning. PNAS. 2015;112:7159–64.

36. Roscoe BP, Bolon DNA. Systematic exploration of ubiquitin sequence, E1 activation efffciency, and experimental fitness in yeast. J Mol Biol. 2014;426:2854–70.

37. Roscoe BP, Thayer KM, Zeldovich KB, Fushman D, Bolon DNA. Analyses of the Effects of All Ubiquitin Point Mutants on Yeast Growth Rate. Journal of Molecular Biology. 2013;425:1363–77.

38. Spencer JM, Zhang X. Deep mutational scanning of S. pyogenes Cas9 reveals important functional domains. Scientific Reports. 2017;7:16836.

39. Starr TN, Greaney AJ, Hilton SK, Ellis D, Crawford KHD, Dingens AS, et al. Deep Mutational Scanning of SARS-CoV-2 Receptor Binding Domain Reveals Constraints on Folding and ACE2 Binding. Cell. 2020;182:1295–1310.e20.

40. Stiffier MA, Hekstra DR, Ranganathan R. Evolvability as a Function of Purifying Selection in TEM-1 β-Lactamase. Cell. 2015;160:882–92.

41. Suiter CC, Moriyama T, Matreyek KA, Yang W, Scaletti ER, Nishii R, et al. Massively parallel variant characterization identifies NUDT15 alleles associated with thiopurine toxicity. Proc Natl Acad Sci U S A. 2020;117:5394–401.

42. Weile J, Sun S, Cote AG, Knapp J, Verby M, Mellor JC, et al. A framework for exhaustively mapping functional missense variants. Molecular Systems Biology. 2017;13:957.

43. Tinberg CE, Khare SD, Dou J, Doyle L, Nelson JW, Schena A, et al. Computational design of ligand-binding proteins with high affinity and selectivity. Nature. 2013;501:212–6.

44. Rollins NJ, Brock KP, Poelwijk FJ, Stifffer MA, Gauthier NP, Sander C, et al. Inferring protein 3D structure from deep mutation scans. Nat Genet. 2019;51:1170–6.

45. Schmiedel JM, Lehner B. Determining protein structures using deep mutagenesis. Nat Genet. 2019;51:1177–86.

46. Ke S, Anquetil V, Zamalloa JR, Maity A, Yang A, Arias MA, et al. Saturation mutagenesis reveals manifold determinants of exon definition. Genome Res. 2018;28:11–24.

47. Kircher M, Xiong C, Martin B, Schubach M, Inoue F, Bell RJA, et al. Saturation mutagenesis of twenty disease-associated regulatory elements at single base-pair resolution. Nat Commun. 2019;10:3583.

48. Melnikov A, Murugan A, Zhang X, Tesileanu T, Wang L, Rogov P, et al. Systematic dissection and optimization of inducible enhancers in human cells using a massively parallel reporter assay. Nature Biotechnology. 2012;30:271–7.

49. Patwardhan RP, Hiatt JB, Witten DM, Kim MJ, Smith RP, May D, et al. Massively parallel functional dissection of mammalian enhancers in vivo. Nature Biotechnology. 2012;30:265–70.

50. Frazer J, Notin P, Dias M, Gomez A, Min JK, Brock K, et al. Disease variant prediction with deep generative models of evolutionary data. Nature. 2021;599:91–5.

51. Gray VE, Hause RJ, Luebeck J, Shendure J, Fowler DM. Quantitative Missense Variant Effect Prediction Using Large-Scale Mutagenesis Data. cels. 2018;6:116–124.e3.

52. Wu Y, Li R, Sun S, Weile J, Roth FP. Improved pathogenicity prediction for rare human missense variants. Am J Hum Genet. 2021;108:1891–906.

53. AVE Alliance Founding Members. The Atlas of Variant Effects (AVE) Alliance: understanding genetic variation at nucleotide resolution [Internet]. Zenodo; 2021 Mar. Available from: https://zenodo.org/record/4989960

54. Impact of Genomic Variation on Function (IGVF) Consortium [Internet]. Genome.gov. [cited 2021 Nov 29]. Available from: https://www.genome.gov/Funded-Programs-Projects/Impact-of-Genomic-Variation-on-Function-Consortium

55. International Common Disease Alliance [Internet]. [cited 2021 Nov 29]. Available from: https://www.icda.bio/

56. Esposito D, Weile J, Shendure J, Starita LM, Papenfuss AT, Roth FP, et al. MaveDB: an open-source platform to distribute and interpret data from multiplexed assays of variant effect. Genome Biology. 2019;20:223.

57. Findlay GM, Boyle EA, Hause RJ, Klein JC, Shendure J. Saturation editing of genomic regions by multiplex homology-directed repair. Nature. 2014;513:120–3.

58. Richards S, Aziz N, Bale S, Bick D, Das S, Gastier-Foster J, et al. Standards and guidelines for the interpretation of sequence variants: a joint consensus recommendation of the American College of Medical Genetics and Genomics and the Association for Molecular Pathology. Genet Med. 2015;17:405–24.

59. Rubin AF, Gelman H, Lucas N, Bajjalieh SM, Papenfuss AT, Speed TP, et al. A statistical framework for analyzing deep mutational scanning data. Genome Biology. 2017;18:150.

60. den Dunnen JT, Dalgleish R, Maglott DR, Hart RK, Greenblatt MS, McGowan-Jordan J, et al. HGVS Recommendations for the Description of Sequence Variants: 2016 Update. Human Mutation. 2016;37:564–9.

61. Hart RK, Rico R, Hare E, Garcia J, Westbrook J, Fusaro VA. A Python package for parsing, validating, mapping and formatting sequence variants using HGVS nomenclature. Bioinformatics. 2015;31:268–70.

62. Wang M, Callenberg KM, Dalgleish R, Fedtsov A, Fox NK, Freeman PJ, et al. hgvs: A Python package for manipulating sequence variants using HGVS nomenclature: 2018 Update. Human Mutation. 2018;39:1803–13.

63. Kluyver T, Ragan-Kelley B Pé, Rez F, Granger B, Bussonnier M, et al. Jupyter Notebooks – a publishing format for reproducible computational workflows. Positioning and Power in Academic Publishing: Players, Agents and Agendas. IOS Press; 2016;87–90.

64. Fowler DM, Araya CL, Gerard W, Fields S. Enrich: software for analysis of protein function by enrichment and depletion of variants. Bioinformatics. 2011;27:3430–1.

